# Amyloid aggregation of *Streptococcus mutans* Cnm influences its collagen-binding activity

**DOI:** 10.1101/2021.06.09.447825

**Authors:** Nicholas de Mojana di Cologna, Sandip Samaddar, Carolina A. Valle, Jonathan Vargas, Alejandro Aviles-Reyes, Joyce Morales, Tridib Ganguly, Roberta Pileggi, L. Jeannine Brady, José A. Lemos, Jacqueline Abranches

**Author notes:** Address correspondence to Jacqueline Abranches.

## Abstract

The glycosylated collagen- and laminin-binding surface adhesin Cnm is present in approximately 20% of *S. mutans* clinical isolates and is associated with systemic infections and increased caries risk. Other surface-associated collagen-binding proteins of *S. mutans* such as P1 and WapA have been demonstrated to form an amyloid quaternary structure with functional implications within biofilms. *In silico* analysis predicted that the β-sheet rich N-terminal collagen-binding domain (CBD) of Cnm has propensity for amyloid aggregation, whereas the threonine-rich C-terminal domain was predicted to be disorganized. In this study, thioflavin-T fluorescence and electron microscopy were used to show that Cnm forms amyloids either in its native glycosylated or recombinant non-glycosylated forms and that the CBD of Cnm is the main amyloidogenic unit of Cnm. We then performed a series of *in vitro*, *ex vivo* and *in vivo* assays to characterize the amylogenic properties of Cnm. In addition, Congo red birefringence indicated that Cnm is a major amyloidogenic protein of *S. mutans* biofilms. Competitive binding assays using collagen-coated microtiter plates and dental roots, a substrate rich in collagen, revealed that Cnm monomers inhibit *S. mutans* binding to collagenous substrates whereas Cnm amyloid aggregates lose this property. Thus, while Cnm contributes to recognition and initial binding of *S. mutans* to collagen-rich surfaces, Cnm amyloid aggregation appears to represent a mechanism to modulate this activity in mature biofilms.

**IMPORTANCE:** *Streptococcus mutans* is a keystone pathogen that promotes caries by acidifying the dental biofilm milieu. The collagen- and laminin-binding glycoprotein Cnm is a virulence factor found in about 20% of the clinical isolates of *S. mutans*. Expression of Cnm by *S. mutans* is associated with niche expansion, allowing colonization of multiple sites in the body including collagen-rich surfaces such as dentin and heart valves. Here, we demonstrate for the first time that Cnm function appears to be modulated by its aggregation status. As a monomer, its primary function is to promote attachment to collagenous substrates via its collagen binding domain (CBD). However, in later stages of biofilm maturation, the same CBD of Cnm self-assembles into amyloid fibrils, losing the ability to bind to collagen and likely becoming a component of the biofilm matrix. Our findings shed light into the role of functional amyloids in *S. mutans* pathobiology and ecology.

## INTRODUCTION

Amyloids represent an evolutionarily conserved fibrillar, cross-β sheet quaternary structure of certain proteins in which the β-sheets laterally assemble in a non-covalent polymer to form fibers (1–4). Amyloid aggregates typically form structures with diameters ranging from 6-12 nm and share common biophysical properties such as detergent and protease resistance, sensitivity to formic acid dissolution, uptake of amyloidophilic dyes, and birefringent properties when stained with Congo red (4, 5). Although amyloids are commonly associated with human diseases related to protein misfolding, functional amyloids are known to play important biological roles across all domains of life (6). In addition to polysaccharides and extracellular DNA (eDNA), amyloids are increasingly detected as constituents of extracellular biofilm matrices where they can influence growth at air-water interfaces (7), serve as a functional sink for quorum sensing molecules (8), or act as a scaffold to help maintain 3D architecture allowing for the formation of channels and wrinkles that determine spatial distribution of species within multi-species biofilms (1, 3, 9–13). Other functional roles of amyloids include regulation of genes and cell fate, toxicity against other microorganisms and host cells, and immune regulation of host responses (12). Several bacterial functional amyloids have been described in both Gram-positive and Gram-negative organisms, including the curli proteins from *Escherichia coli* and *Salmonella spp*, *Bacillus subtilis* TasA, *Pseudomonas aeruginosa* FapBC*, Staphylococcus aureus* Bap and Phenol-Soluble Modulins (PSM), *Enterococcus faecalis* Esp, and *Streptococcus mutans* P1 (also known as SpaP or Antigen I/II), WapA and SMU_63c (12, 14–19). Multiple factors including pH, salt concentration and surface hydrophobicity have been reported to influence the propensity of proteins to form amyloids within bacterial biofilms (1, 10, 15, 20, 21).

A resident of the dental biofilm, *S. mutans* is a keystone pathogen in dental caries. The cariogenic potential of *S. mutans* can be largely attributed to several characteristics that are hallmarks of its successful coevolution with humans. First, *S. mutans* genomes encode a number of glucosyltransferase enzymes that convert sucrose into large glucan polymers, which comprise the bulk of the extracellular polymeric matrix that are critical for the establishment and maintenance of the dental biofilm (22). In addition, *S. mutans* thrives in carbohydrate-rich environments as it can metabolize a wide range of carbohydrates, generating acids (acidogenicity) and surviving and continuing to undergo glycolysis even at pHs below 5.0 (aciduricity) (23, 24).

In addition to the so-called sucrose-dependent colonization mechanism that is based on glucan polymer synthesis, *S. mutans* also possesses sucrose-independent colonization mechanisms whereby cell surface-associated adhesins such as P1 and WapA mediate attachment to the salivary pellicle that coats the tooth enamel (25, 26). Of note, P1 and WapA also mediate binding to collagenous-rich surfaces (27). More recently, two closely-related surface-associated collagen-binding proteins (CBPs), Cnm and Cbm, were identified in approximately 20% of *S. mutans* strains (28–30). Cnm was shown to mediate invasion of epithelial and oral cell lines and binding to collagenous and laminin-rich substrates present in oral (dentin, cementum and roots) and extra-oral environments (heart valves) (27, 31–33). Cnm and Cbm were shown to be important virulence factors implicated in systemic infections such as infective endocarditis, cerebral microbleeds and hemorrhagic stroke (34–38). Moreover, clinical studies have linked oral infection with CBP^+^ *S. mutans* strains with increased caries risk and poor caries outcomes (23, 27, 39, 40).

While Cnm has a predicted molecular mass of 54 kDa, the Cnm protein migrates at approximately 120 kDa when separated by SDS-PAGE. Recently, we discovered that this aberrant migration was, in great part, a result of Cnm post-translational modification through glycosylation. Specifically, we identified a novel protein glycosylation machinery *pgf* (<u>p</u>rotein <u>g</u>lycosyltransferase) in *S. mutans* (41) that was co-transcribed with *cnm* and responsible for its glycosylation in serotype *f* OMZ175 strain (42). Inactivation of *pgf* genes resulted in a Cnm protein that migrated at approximately 90 kDa and that is highly susceptible to proteolysis, indicating a critical role for glycosylation in Cnm function and stability (41, 42). *In silico* analysis revealed that the threonine-rich repeat domain located in the Cnm C-terminus is subjected to glycosylation, while the β-sheet rich N-terminus where the conserved collagen binding domain (CBD) resides is not modified by the Pgf glycosylation machinery (42). In addition, the Pgf system was shown to also modify WapA, whose N-terminal truncation product Antigen A (AgA) was shown to have amyloidogenic properties (41). Interestingly, homology models revealed a high similarity between the tertiary structures of the Cnm CBD with that of AgA (27). Based on the similarity between AgA and the Cnm CBD, we hypothesized that Cnm may also self-assemble into amyloid fibers. In the present study, we confirmed this prediction by using a panel of wild-type and mutant strains and different variant forms of purified Cnm (native glycosylated, recombinant non-glycosylated and recombinant CBD). We show that Cnm is a major amyloidogenic protein in the Cnm^+^ *S. mutans* strain OMZ175, with its CBD being sufficient for amyloidogenesis but possibly requiring the threonine-rich repeat for maintenance of stability, especially in acidic environments. We also demonstrate that when in monomeric form, the main function of Cnm is collagen binding. However, when Cnm self-assembles into amyloids, its ability to competitively inhibit binding of *S. mutans* cells to collagenous substrates is lost. Hence amyloid formation by Cnm likely acts as a negative regulatory mechanism to modulate collagen-binding activity within *S. mutans* biofilms.

## RESULTS

### *In silico* analyses suggests that the collagen binding domain (CBD) of Cnm is aggregation-prone

To evaluate the potential for Cnm amyloid formation, we used the bioinformatics tool AmyloPred2 to analyze the primary structure of Cnm from *S. mutans* OMZ175. The analysis predicted a high frequency of aggregation-prone amino acid residues in the N-terminal portion (within the CBD) (Figure 1A and 1B), which coincides with the predicted β-sheet-rich region modelled using Swiss-Model (Figure 1C). The threonine-rich repeat domain of Cnm could not be modeled due to its high degree of structural disorder (defined as lack of fixed three-dimension structure), and therefore can be classified as an intrinsically disordered region (Figure 2A) (11). It is possible, however, that this level of structural disorder is different when the protein is glycosylated. Such a disorder analysis using MobiDB requires a deposited UniProtKB sequence, and for this analysis the Cnm sequence of strain TW871 (UniProtKB C4B6T3) that shares 98% and 100% identity with full-length Cnm and the Cnm CBD from *S. mutans* OMZ175, respectively, was used. To gain further insight into how the aggregation-prone N-terminal domain of Cnm may lead to amyloid formation, we performed homology modeling of a Cnm dimer based on the solved structure of the dimer of Clumping Factor B (ClfB) from *S. aureus* (PDB: 3AU0) on GalaxyHomomer. ClfB was identified by the software as the most structurally similar sequence deposited in the Protein Data Bank. The analysis predicted an interaction between the β-sheet regions of CBDs from different Cnm monomers (Figure 2B). Portions of the β-sheet region of the interacting monomers would also remain available for potential coupling with additional Cnm monomers, while the predicted unstructured acidic and glycosylated threonine-rich domains would protrude outwards without active participation in the predicted interaction (Figure 2B).

**FIG 1.**
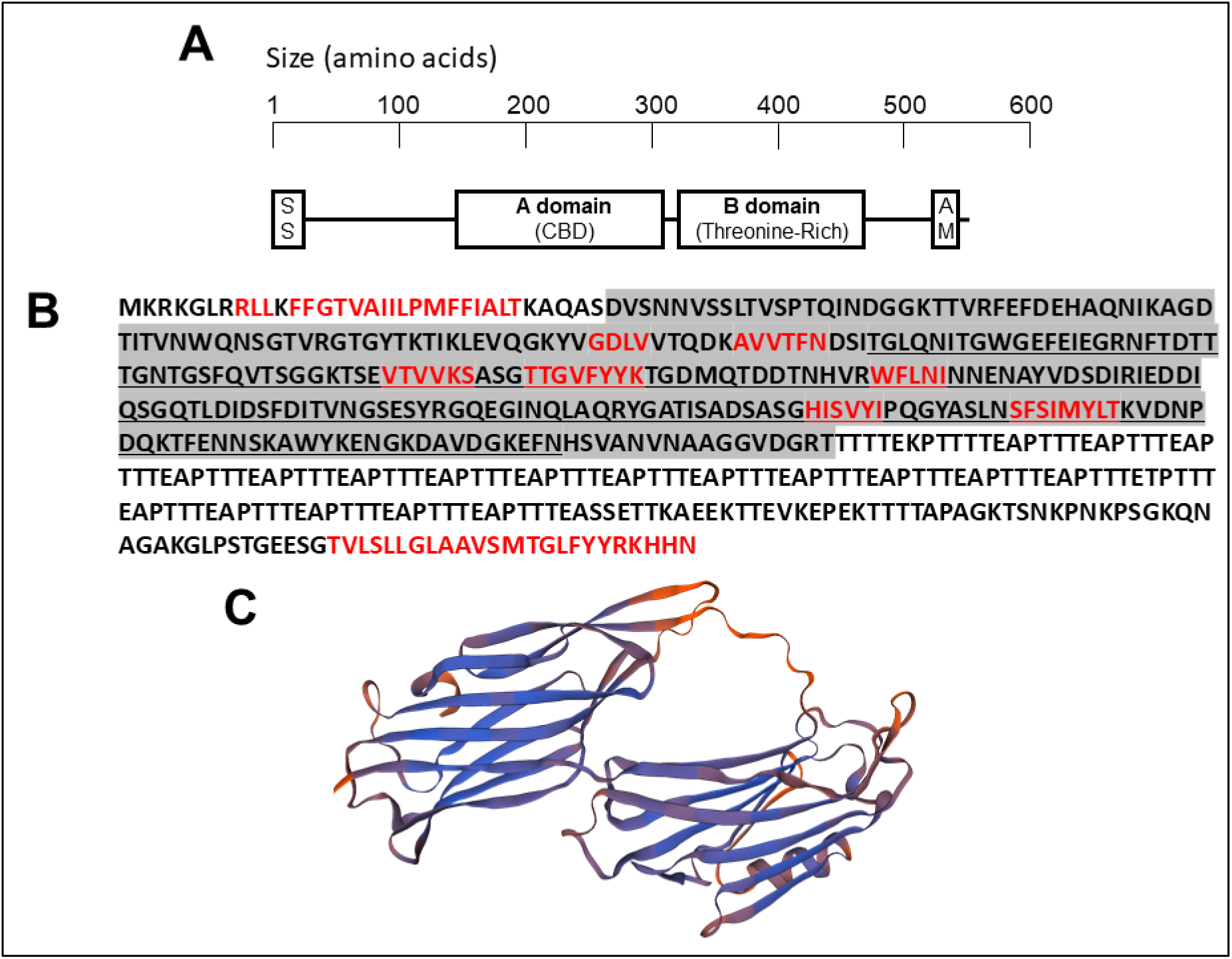
The β-sheet-rich Collagen-Binding Domain of *S. mutans* Cnm is predicted to be amyloidogenic. (A) Schematic representation of Cnm, including the Secretion Signal (SS), the A and B domains, and the LPXTG Anchoring Motif (AM). (B) Aggregation-prone peptides predicted by AmylPred2 are shown in red. The CBD is underlined, and the portion of the sequence that was modeled by Swiss-Model is highlighted in gray. (C) Swiss-Model homology model of the CBD of Cnm from *S. mutans* OMZ175 (based on model SMTL ID 2f6a.1).

**FIG 2.**
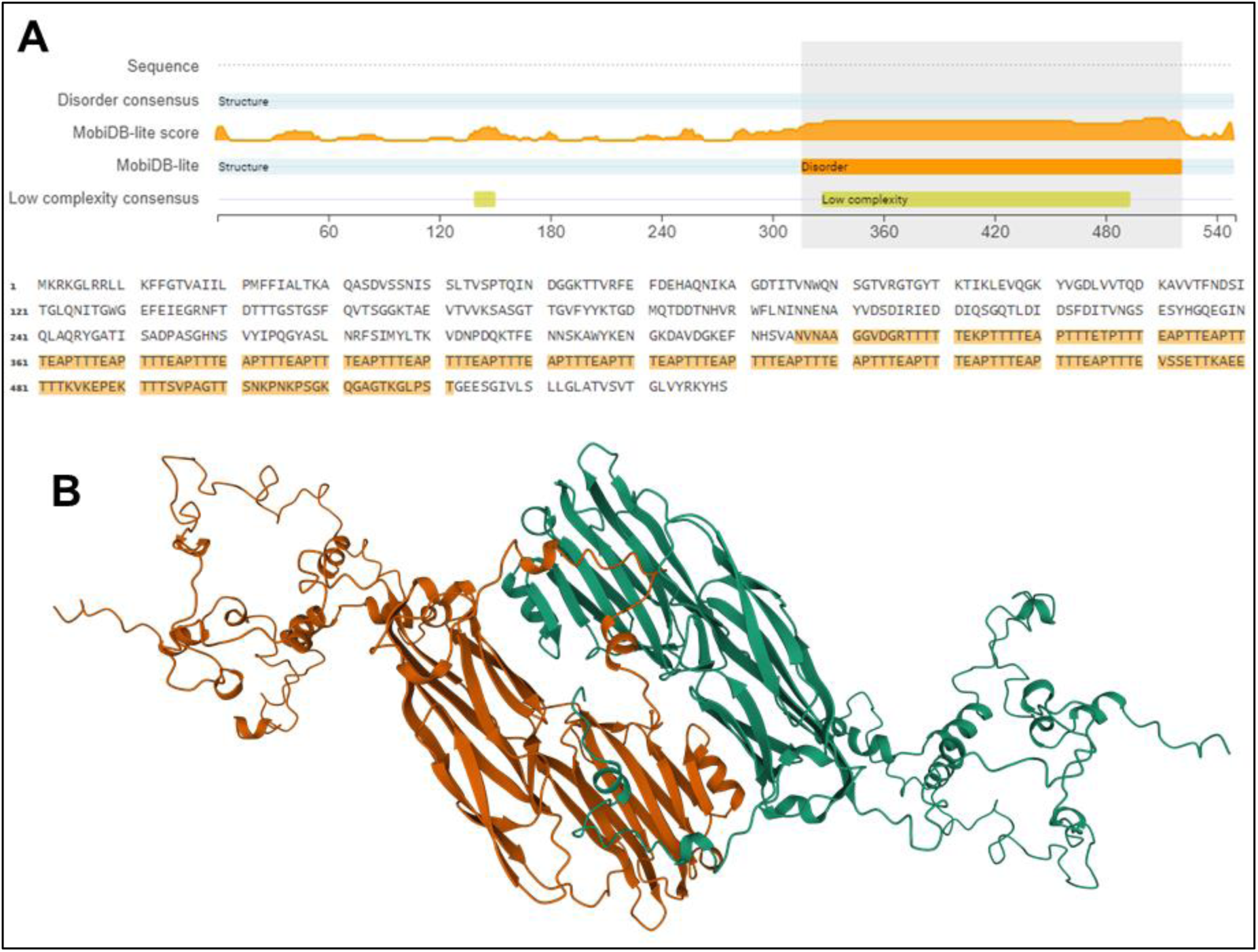
The Cnm threonine-rich B domain as an intrinsically disordered region. (A) MobiDB estimation of structural disorder highlights a lack of fixed 3D structure of the B domain of the deposited Cnm sequence UniProtKB C4B6T3. (B) GalaxyHomomer modeling of full-length Cnm, based on homology with the *S. aureus* ClfB dimer, combines ordered and disordered regions and illustrates likely interactions between β-sheet-rich CBDs of two different Cnm monomers (each in a different color, orange or green). The acidic and possibly glycosylated threonine-rich domains protrude outwards.

### Cnm and its variant forms uptake the amyloidophilic dye Thioflavin-T

After *in silico* analyses predicting that Cnm is prone to aggregation, we performed ThT fluorescence assays using purified Cnm in the presence or absence of the amyloid inhibitor tannic acid (43). In addition to full-length glycosylated Cnm purified from *S. mutans* OMZ175, we also tested a recombinant full-length Cnm (rCnm) and a truncated Cnm containing the CBD only (rCBD) purified from *E. coli*. Of note, both recombinant protein versions are not glycosylated as *E. coli* lacks the Pgf machinery required for Cnm glycosylation. As shown in Figure 3A, all three tested forms of Cnm incorporate the amyloidophilic ThT dye when amyloidogenesis was stimulated by stirring in pH 7.4. Mechanical agitation of purified proteins is a common strategy to promote nucleation, enhancing amyloid fibril assembly (44). Moreover, ThT incorporation by all three Cnm versions was significantly inhibited by the amyloid inhibitor Tannic Acid (Fig. 3A). Collectively, these results suggest that both glycosylated and non-glycosylated Cnm can form amyloids and that the CBD is the main amyloidogenic unit of Cnm.

**FIG 3.**
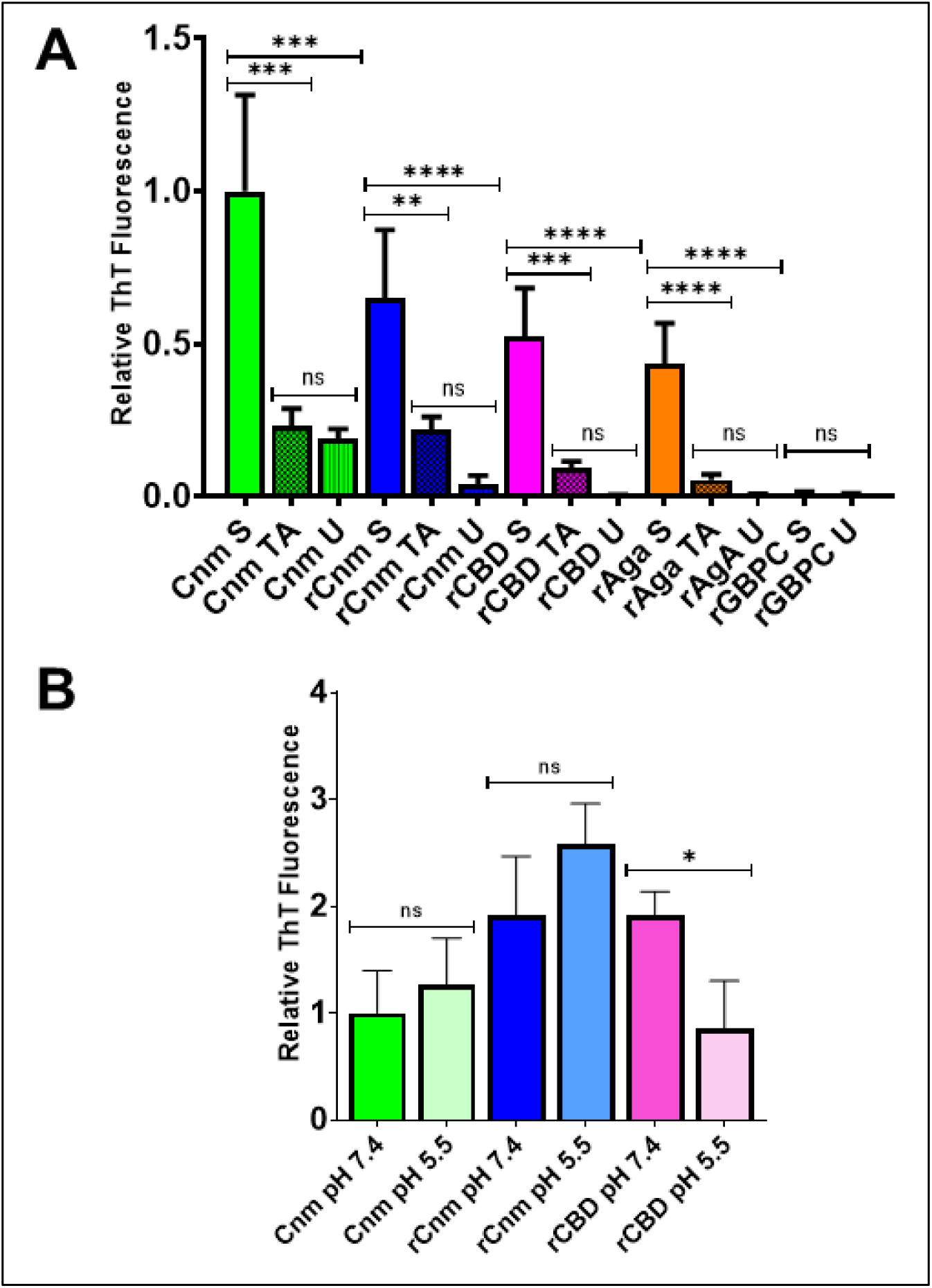
Thioflavin-T (ThT) fluorescence assay of native glycosylated *S. mutans* Cnm, and recombinant non-glycosylated Cnm (rCnm and rCBD) purified from *E. coli*. (A) ThT uptake by purified Cnm. rAgA and rGbpC were used as a positive and negative controls, respectively. S: Stirred; TA: Stirred with Tannic Acid; U: Unstirred. All values were normalized by molarity and relative to stirred Cnm (B) Effect of pH on amyloidogenic properties of native Cnm and derivatives. All values were normalized by molarity and are relative to Cnm at pH 7.4. Experiments were performed at least in triplicate with statistical analyses performed using one-way ANOVA (for each group shown in Panel A), or nonparametric T tests (for each pair shown in Panel B). ns = not significant; * = p<0.05; ** = p<0.01; *** = p<0.001; **** = p<0.0001.

To determine whether pH can influence Cnm fibril assembly, we performed the same ThT assay under a neutral pH (7.4) and an acidic pH (5.5), the latter being similar to the pH of enamel dissolution provoked during dental plaque acidification by *S. mutans* (Figure 3B). While the full-length forms of Cnm (Cnm and rCnm) did not show significant differences in dye uptake based on pH, ThT uptake by rCBD was significantly decreased at pH 5.5 when compared to pH 7.4 (p<0.05). This suggests that the unstructured threonine-rich domain may contribute to quaternary structure stability at lower pH.

### Transmission electron microscopy confirms that *S. mutans* Cnm and its variants form amyloids

A defining feature of amyloid aggregates is protease resistance (45–47). In *S. mutans,* protein amyloids form mat-like composite aggregates in which typical amyloid fibrillar structures are only revealed following protease digestion of protein monomers or protease-sensitive oligomeric intermediates (48). To obtain direct evidence that Cnm forms amyloids, stirred samples of full-length glycosylated Cnm, non-glycosylated Cnm (rCnm) and rCBD were treated with proteinase K and visualized by transmission electron microscopy (TEM) (Figure 4). Fibrillar material indicative of amyloid aggregates was visualized for the recombinant forms of non-glycosylated Cnm (rCnm and rCBD). In contrast, a mat-like amyloid structure was visualized for glycosylated native Cnm (Figure 4). Because *S. mutans* amyloid mats have been confirmed to exhibit the same characteristic X-ray fiber diffraction pattern as isolated amyloid fibers (48), and because glycosylation of Cnm has been shown to protect this protein from proteolytic degradation (41), the visualization of amyloid mats rather than fibers by TEM confirms the formation of amyloid by native glycosylated Cnm.

**FIG 4.**
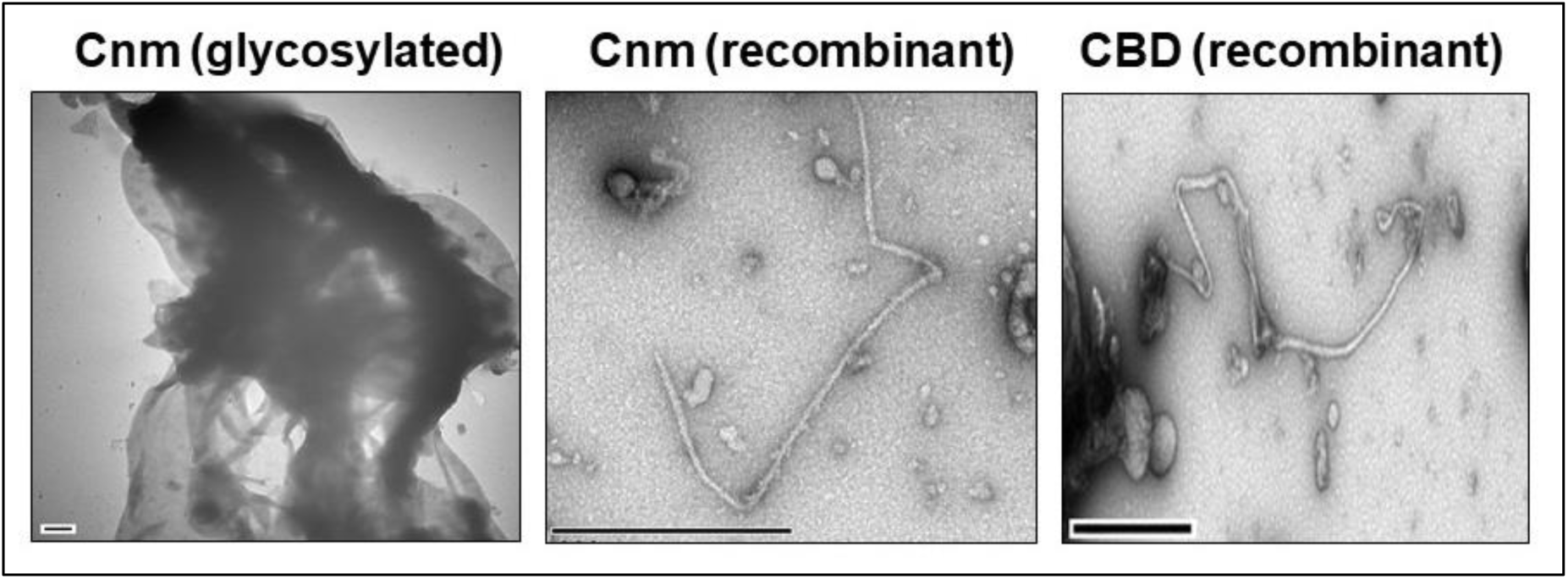
Transmission electron microscopy of stirred *S. mutans* Cnm, rCnm and rCBD following treatment with proteinase K. Purified proteins were stirred to induce amyloid formation, then treated with proteinase K to remove residual protease-sensitive monomers. Fibrillar material was visualized in the non-glycosylated rCnm and rCBD preparations, whereas the native glycosylated Cnm sample displayed a mat-like structure, possibly due to proteinase K resistance. Scale bars = 500 nm.

### Cnm is a major amyloidogenic protein in OMZ175 biofilms

Congo red is an amyloidophilic dye which provides amyloid-containing cell foci an aberrant coloration, typically yellow or orange-green, when viewed with polarized light through crossed filters (49). To determine whether Cnm amyloid fibrils can be detected in the *S. mutans* OMZ175 biofilm matrix, Congo red birefringence assays were performed on mature (5 days-old) biofilms using a panel of strains that include the parent OMZ175, the *cnm* deletion mutant (OMZ175Δ*cnm*) and the *srtA* deletion mutant (OMZ175Δ*srtA*). The *srtA* gene codes for the sortase A enzyme such that the OMZ175Δ*srtA* cannot cross-link LPXTG proteins, like Cnm, to the cell envelope. Moreover, previous work has shown that biofilm-associated amyloid formation was diminished in an *S. mutans* Δ*srtA* strain (16). As shown in Figure 5A, strongly birefringent, tightly organized amyloid-containing aggregates were observed in mature biofilms formed by OMZ175, while less intense, more diffuse staining was observed in the Δ*cnm* and Δ*srtA* strains grown in media containing glucose (Figure 5A). Low magnification visualization (data not shown) and quantification of amyloid aggregates per slide revealed that OMZ175 biofilms display significantly more birefringent foci than either mutant strain, with no significant differences between the two mutants (Figure 5B).

**FIG 5.**
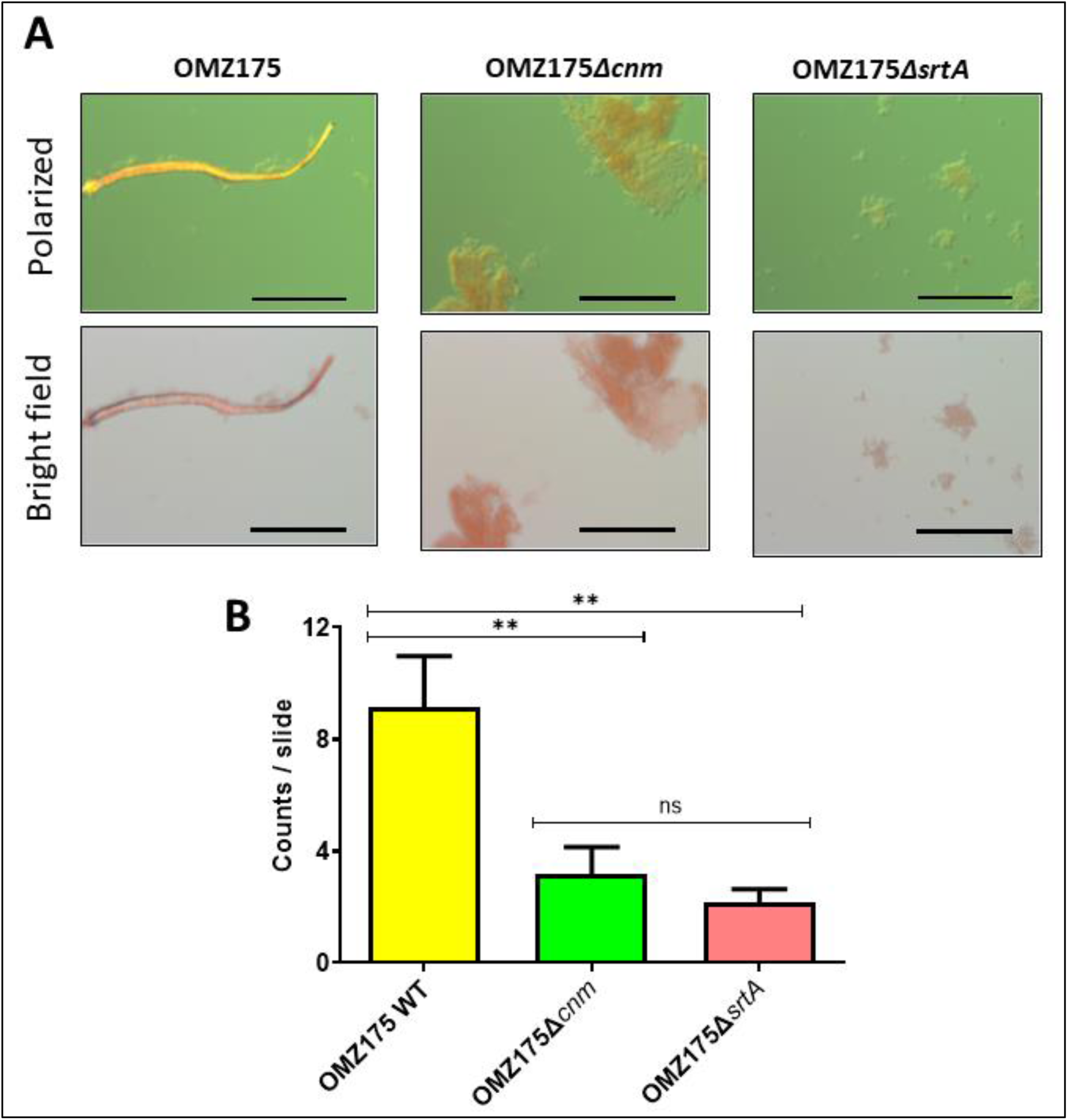
Cnm is a major amyloidogenic protein of *S. mutans* OMZ175 biofilms. Five-day old biofilm biomass of OMZ175, OMZ175Δ*cnm* and OMZ175Δ*srtA* strains were stained with Congo red and visualized by polarizing light microscopy with crossed filters. (A) Representative micrographs of biofilm masses as seen under bright field and cross-polarized light filters. Scale bars = 100 µm. (B) Counts of birefringent foci per slide for each strain confirm that Cnm amyloids are prevalent within *S. mutans* OMZ175 biofilms when compared to OMZ175Δ*cnm* and OMZ175Δ*srtA* strains (p= 0.0297). Experiments were performed at least in triplicate with statistical analyses performed using one-way ANOVA: ** = p<0.01; ns = non-significant.

### CBD monomers, but not amyloid fibers, competitively inhibit cell binding to collagenous substrates

To gain insight into the biological relevance of the amyloidogenic property of Cnm, we developed a competitive binding assay in which purified rCBD monomers or fibrils were tested for their ability to interfere with the ability of the OMZ175 strain to adhere to collagen-coated microtiter plate wells or to premolar roots of which collagen is the main organic component (50, 51). We used rCBD for these assays because it can be produced in large quantities and encompasses the domain that mediates collagen binding and can form amyloid fibers. Figure 6A shows that rCBD monomers inhibited OMZ175 binding to immobilized collagen whereas rCBD-derived amyloid fibrils did not (p<0.0001). These results indicate that the rCBD monomers, but not the amyloid aggregates, can bind to collagen-rich surfaces thereby making it less available for binding by OMZ175 cells. In an *ex vivo* model, the competitive binding assay was performed using saliva-coated dental root sections, with significant similar results obtained (Figure 6B). These results are consistent with *in silico* prediction of Cnm oligomerization via the β-sheets of the CBD domains (Figure 2B), in that once Cnm-Cnm interactions occur the CBD should no longer be available for interaction with collagen.

**FIG 6.**
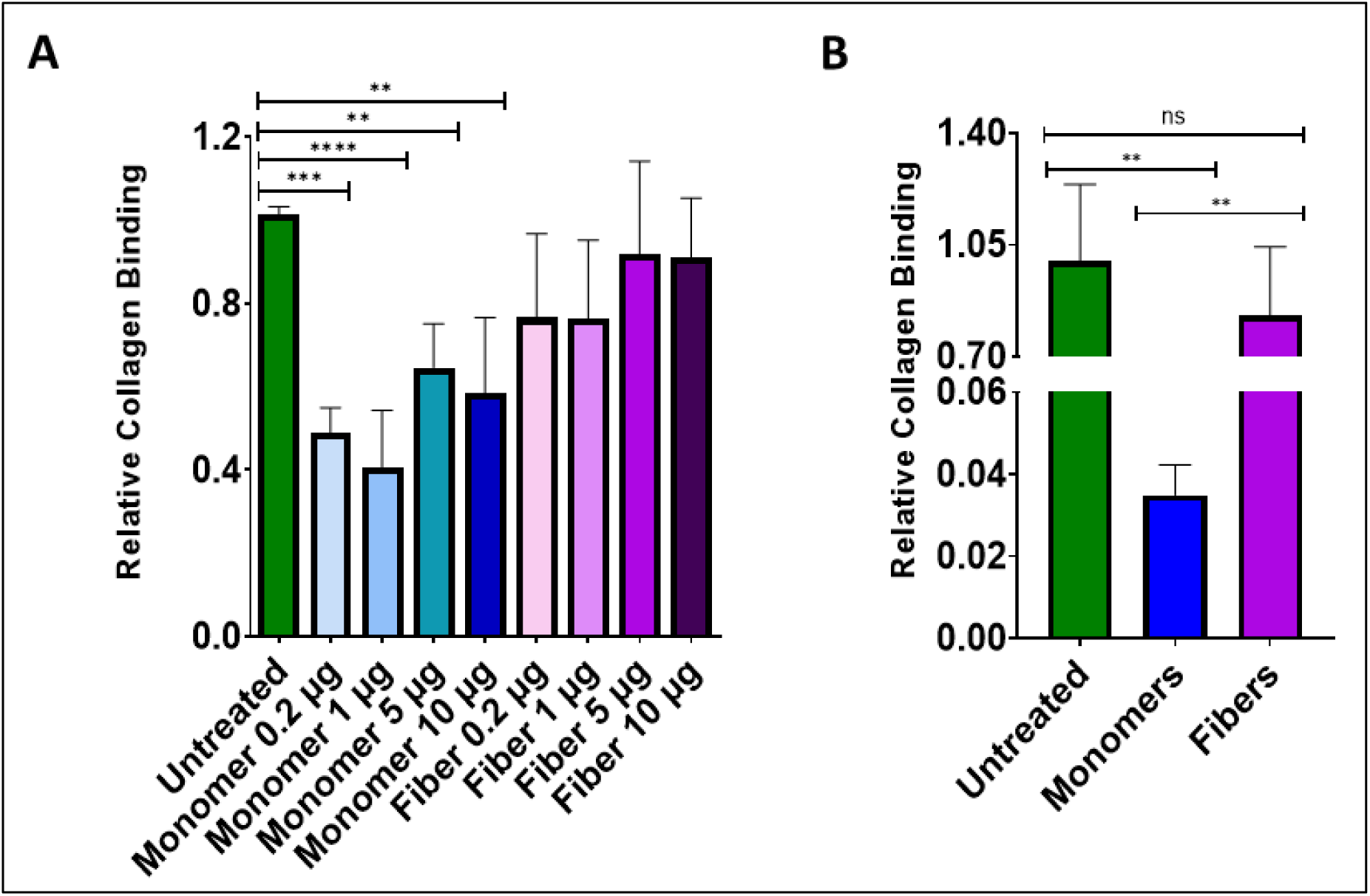
Amyloid aggregation of *S. mutans* Cnm diminishes its binding to collagenous substrates. (A) *In vitro* competitive inhibition assay in which collagen-coated microtiter plates were pre-incubated with increasing concentrations of rCBD monomers, or corresponding amyloid fibrils, prior to incubation with OMZ175 cells. Values indicate binding relative to untreated wells that did not receive rCBD monomers or fibrils. rCBD monomers, but not amyloid fibrils, significantly inhibited OMZ175 adherence. (B) *Ex vivo* competitive inhibition assay using pre-molar roots. Pre-incubation with 20 µg rCBD monomers, but not with 20 µg of amyloid fibers, significantly diminished binding of OMZ175 to the root sections, resulting in lowered CFU recovery. Values are relative to untreated (no monomers or fibrils added). Experiments were performed at least in triplicate with statistical analyses performed using one-way ANOVA: (**) p<0.01; (***) p<0.001; (****) p<0.0001.

## DISCUSSION

Microbial amyloids have been a focus of increased attention due to their wide array of biofilm-associated functions, which include organization of biofilm structure, modulation of biofilm hydrophobicity and mechanical properties, control of quorum sensing, and regulation of genes, toxicity and modulation of host response (1, 8–13, 52). Functional amyloids have been discovered in pathogenic and non-pathogenic bacteria, and their formation is generally associated with growth in biofilms (1, 10, 45). The discovery of multiple functional amyloids in *S. mutans* (WapA, P1 and SMU_63c) made us wonder if Cnm, a surface protein that is also rich in β-sheet structure, could be part of this group. Here, we show, for the first time, that Cnm function is modulated by its aggregation status.

An amyloid is a non-covalent oligomer of extended inter-molecularly hydrogen-bonded β-sheets that self-assemble to form fibers. Regardless of the origin, amyloids share several properties, which includes a fibrillar structure when viewed by electron microscopy, enhanced birefringence on binding to Congo red, and a cross-β sheet structure in which the β-strands run perpendicular to the fiber axis resulting in a characteristic X-ray fiber diffraction pattern (53). Our *in silico* analyses suggested that the CBD portion of Cnm assembles into amyloid aggregates, whereas the glycosylated threonine-rich repeat likely does not contribute to the amyloid fibril assembly. During oligomerization, the β-sheet structures of the monomeric units would interact, with the glycosylated threonine-rich domains protruding outwards. This model is consistent with our *in vivo* and *ex vivo* collagen-binding competitive inhibition assays, in which we observed that purified rCBD monomers competed with intact cells of *S. mutans* for binding to collagen-rich surfaces while rCBD-derived amyloid did not. In other words, loss of the collagen-binding function of Cnm following amyloid aggregation suggests that the same β-sheet rich region that mediates collagen binding also functions as the elongation point for fibril formation, thus rendering it unavailable for its primary adhesive function. Depending on the protein, formation of amyloids can result in a loss or a gain of function of the aggregating polypeptide (4). Hence, amyloid aggregation represents a potential mechanism to regulate protein function as needed in different environments (54, 55). The oligomerization of Cnm into amyloids does not seem to be dependent upon glycosylation by the Pgf machinery, since native glycosylated and non-glycosylated recombinant proteins both activated ThT fluorescence and displayed visible amyloid fibers when visualized by microscopy. The contribution of glycosylation to amyloidogenic proteins is still a relatively unexplored field, although the effect of glycosylation of the Amyloid Precursor Protein of Aβ in Alzheimer’s disease has gained attention due to its regulatory role in the protein’s proteolytic processing (56, 57). Bacterial amyloids derived from glycoproteins have now been reported as components of the structural extracellular matrix of biofilms rich in ammonia-oxidizing bacteria and nitrite-oxidizing bacteria grown in aerobic and granular sludge (58). Further characterization of glycoprotein-derived amyloids is warranted as these may demonstrate common features in different domains of life.

In addition to its correlation with increased caries risk, *S. mutans* Cnm has also been shown to contribute to increased acid tolerance (59). Therefore, it was relevant to investigate if the newfound amyloidogenic properties of Cnm are modulated by acidic pH. Based on ThT assays of Cnm, rCnm and rCBD stirred in neutral compared to acidic pH, we found that in contrast to native and rCnm, the rCBD variant forms fewer amyloid fibrils at pH 5.5 than at pH 7.4. This result suggests that the threonine-rich domain may contribute to fiber integrity of full-length Cnm at acidic pH. This effect may be related to the presence of an external layer of acidic threonines to accept protons, thus protecting the β-sheet core from unfolding by charge repulsion (60). This observation strengthens the notion that, although the threonine-rich domain is not related to the collagen-binding activity of Cnm, it supports that activity by contributing to the stability of the Cnm monomer. For instance, a previous study from our group demonstrated that glycosylation of the threonine-rich domain by the Pgf machinery protects Cnm against the proteolytic activity of proteinase K (41). Also, the effect of acidity in promoting amyloidogenesis has been well characterized for Aβ oligomers related to Alzheimer’s disease as a key factor for pathogenicity (61, 62). In *S. mutans*, neutral pH was shown to favor amyloid formation by WapA and P1, while the secreted amyloidogenic protein Smu63c, which appears to be a negative regulator of biofilm cell density and genetic competence, is triggered to assemble into amyloid by acidic pH (15). Of note, formation of *S. aureus* Bap amyloids are also favored at acidic pH (15, 63).

TEM visualization of amyloid material induced by mechanical agitation of purified proteins, followed by treatment with proteinase K, revealed a mat-like structure for glycosylated Cnm but a fibrillar morphology for non-glycosylated Cnm and CBD. Mat-like aggregates have been proposed as a more biologically germane form of *S. mutans* amyloids that represent a supramolecular structure composed of amyloid fibers, monomers, and oligomer intermediates (48) where pure fibers, achieved by proteolytic digestion of *S. mutans* amyloid-containing mats, are likely only seen in laboratory settings. It has also been shown that the addition of monomeric protein to purified fibrils shifts the morphology of fibrils back to mats (48). Amyloid mats derived from purified *S. mutans* C123 (the amyloid forming truncation derivative of P1), AgA (the amyloid forming truncation product of WapA), and Smu_63c, as well as purified fibers produced by protease treatment of the mats, all demonstrated classical X-ray fiber diffraction patterns with distinct intensities at 4.8 Å in the meridional direction corresponding to the separation of strands in a β-sheet, and distinct equatorial intensities at 10 Å corresponding to the distance between stacked β-sheets (48). Our current results revealed that proteinase K treatment of aggregated native glycosylated Cnm did not disrupt the amyloid mats, confirming our previous finding that glycosylation of Cnm confers resistance to proteinase K degradation of monomers (41). In contrast, protease treated aggregates of non-glycosylated rCnm and rCBD were more fibrillar in nature. The visualization of amyloid mats and fiber morphologies, in conjunction with ThT uptake assays provide direct evidence of amyloid formation by *S. mutans* Cnm.

To confirm Cnm amyloidogenic properties *in vivo*, Congo red-induced birefringence was evaluated for mature biofilms of the wild type, Δ*cnm* and Δ*srtA* strains. Of note, *S. mutans* harbors only one sortase-encoding gene, and inactivation of *srtA* results in a complete lack of LPXTG motif-containing proteins on the cell surface (64), including the amyloidogenic proteins P1, WapA and Cnm. While not every sortase substrate is amyloidogenic (15), most amyloid forming proteins identified in other bacteria are anchored to the cell surface (17, 63, 65). In *S. mutans*, elimination of *srtA* negatively impacted biofilm-associated amyloid formation suggesting that cell surface protein anchoring contributes to this process (16). Birefringent foci were visualized in 5 days-old *S. mutans* OMZ175 biofilms, and surprisingly, the OMZ175Δ*cnm* mutant showed similarly diminished birefringent properties as the OMZ175Δ*srtA* mutant compared to the parent OMZ175 strain. This indicates that Cnm is the main amyloidogenic protein of strain OMZ175 and is present in amyloid form within mature biofilms likely as a component of the biofilm matrix. Notably, while *spaP* (encoding P1) is present in OMZ175 genome, the expression of P1 appears to be down-regulated in Cnm^+^ strains (66), hence the pronounced impact of *cnm* deletion on detection of biofilm-associated amyloids.

A shift in function between monomeric and fibrillar forms of an amyloidogenic protein has been previously observed for *S. aureus* PSMs (19). Monomeric (soluble) and amyloid fibers (insoluble) of PSMα peptides were assessed for their ability to modulate the antibiotic tolerance to ciprofloxacin in *S. aureus* cultures. The presence of PSMα monomers led to a reduction in the number of persister cells tolerant to ciprofloxacin, while the presence of PSMα amyloid fibrils did not alter the persister cell phenotype under the same conditions. It was proposed that the PSMα fibrils might act as reservoirs of the active form of this virulence factor that could be later mobilized if needed during changes in environmental conditions such as pH fluctuation, a factor known to influence fiber formation and dissociation of certain amyloids (61, 67). Another functional shift was also described between monomeric and fibrillar forms of PSM whereby several PSM monomers (PSMα, PSMβ and δ-toxin), known to contribute to biofilm detachment (68), displayed the opposite behavior of promoting biofilm integrity when sequestered into amyloids (18). Based on our collagen-binding competition assays, we observed that Cnm also undergoes a functional shift when in amyloid form. In biofilms of common laboratory strains of *S. mutans* that lack Cnm, amyloid material is more readily detected as the biofilm matures, usually after 60 hours of growth (48). Whether this represents time and concentration-dependent amyloid nucleation leading to polymerization and elongation (69), or depends on other changing environmental conditions within the aging biofilm, is not yet understood. The loss of Cnm collagen-binding activity as the protein transitions to amyloid form strongly suggests that it serves as mechanism to regulate this property once adhesion and colonization have occurred and the biofilm is established. It is possible that Cnm amyloid fibrils present with other macromolecules such as eDNA and polysaccharides in mature biofilms serve a scaffolding function to integrate biofilm cells and provide a substrate for attachment of other members of the biofilm community. Alternatively, accumulation of amyloid material in the matrix may result in consolidation and detachment of mature biofilms, as surface-anchored monomers stop interacting with the collagenous substrates and interact instead with other monomers to form amyloid fibers within the biofilm. Studies are underway to determine the contribution of Cnm and other amyloids to the 3D architecture during progression of *S. mutans* biofilm development. It will be important to establish whether the amyloid form of Cnm alters other functions already described for this virulence factor including oral colonization, and epithelial and endothelial cell invasion (27, 34). Our results provide the foundation to address these questions in future studies. Our findings also shed light on an emerging paradigm of Gram-positive multi-functional proteins and their structural transitions from monomer to amyloid forms as an energy-effective strategy for biofilm modulation (17, 63, 70). Such new aspects of Cnm pathobiology will inform future studies not only in *S. mutans* but also in a myriad of other organisms implicated in biofilm-related disease processes.

## MATERIAL AND METHODS

### Bacterial strains and growth conditions

Strains used in this study are listed in Table and Table*. Escherichia coli* strains were routinely grown in Luria-Bertani (LB) broth medium at 37°C. When required, 100 μg mL^−1^ ampicillin was added to LB broth or to agar plates. *S. mutans* strains were routinely cultured in Brain Heart Infusion (BHI) medium at 37°C in a humidified 5% CO_2_ atmosphere. When required, 1 mg mL^−1^ kanamycin was added to BHI broth or to agar plates.

### Genetic manipulation of *S. mutans*

A sortase A (*srtA*) null-mutant was created in OMZ175 by using the PCR-ligation strategy (71) to replace the *srtA* gene with a non-polar kanamycin marker (72). The 700 bp region flanking the 5’ of the *srtA* gene was amplified using the primers srtA.1F (5’-TGAGTCGCGATAATGATG-3’) and srtA.1RPstI (5’-GAAACCTGA<u>CTGCAG</u>TTGGTATTC-3’), whereas the 700bp flanking the 3’end of *srtA* was amplified using srtA.2FPstI (5’-GAATACCAA<u>CTGCAG</u>TCAGGTTTC-3’) and srtA.2R (5’-GCAGCGGTTCAACTAACTTCTC-3’). The underlined bases correspond to the PstI restriction site that was included for cloning purposes. After amplification, the two PCR fragments were digested with PstI and ligated to a nonpolar kanamycin resistance gene cassette that was obtained as a PstI fragment. The ligation mixture was used to transform *S. mutans* OMZ175, followed by plating onto BHI containing kanamycin (1 mg mL^−1^). The insertional inactivation of *srtA* was confirmed by PCR sequencing.

### *In silico* analysis

The AmylPred2 software (http://aias.biol.uoa.gr/AMYLPRED2/) (73), which employs a consensus of several methods to predict amyloid formation propensity, was used to analyze the primary amino acid sequence of Cnm from *S. mutans* strain OMZ175. Homology modeling of the Cnm monomer was performed with Swiss-Model (74) based on model SMTL ID 2f6a.1, and a homo-oligomerization homology model was obtained based on the *Staphylococcus aureus* Clumping Factor B dimer structure on GalaxyHomomer (75). Protein structure disorder estimations were calculated using MobiDB based on the similar deposited Cnm sequence UniProtKB C4B6T3 (76).

### Expression and purification of recombinant Cnm and CBD

Both recombinant CBD and full-length Cnm (rCBD and rCnm, respectively) were expressed in *E. coli* (41, 77). The *E. coli* strain harboring the pET16b-rCnm plasmid was grown in LB broth containing ampicillin to an optical density at 600 nm of ≈0.5 (37°C and 150 rpm). The expression of the His-tagged protein was induced by the addition of 0.5 mM isopropyl-β-d-thiogalactopyranoside (IPTG) for 20 hours at 24°C. The strain harboring the pET16b-rCBD plasmid was also grown to an optical density at 600 nm of ≈0.5 (37°C and 150 rpm) and induced with 0.5 mM IPTG for 4 hours at 37°C. Cells were lysed and recombinant proteins were purified under native conditions using the Ni-NTA Fast Start Kit (Qiagen) following the manufacturer’s instructions. Halt Protease Inhibitor Single-Use Cocktail (Thermo Fischer Scientific) was added to the lysis buffer as the only modification to the protocol.

### Expression and purification of native Cnm

Native glycosylated Cnm was purified using a custom affinity chromatography column coupled to an anti-rCBD antibody as previously described by our lab (42, 77). Briefly, the OMZ175*ΔvicK* strain, known to overexpresses Cnm (78), was grown overnight and then lysed by bead beating in the presence of Halt Protease Inhibitor Single-Use Cocktail (Thermo Fischer Scientific) and Ethylenediaminetetraacetic acid (EDTA). The soluble protein fraction was bound to a custom NHS–anti-rCBD column with overnight rocking at 4°C. The column was then washed with 15 mL of 1x Phosphate Buffer Saline (PBS) at pH 7.2, and Cnm was eluted using 0.1 M Glycine Buffer (pH 2.5) for 5 minutes. Elutions were immediately neutralized with 1/10 volume of a basic 1 M Tris buffer (pH 8.0). Purified native Cnm was later concentrated and dialyzed against a 100 mM ammonium bicarbonate solution.

### Protein quantification and SDS-PAGE

Protein concentrations were determined with the micro BCA Protein Assay Kit (Thermo Fischer Scientific), using a Bovine Serum Albumin (BSA) standard curve as a reference for linear regression.

Denaturing SDS 10% (m/v) polyacrylamide gels were performed to check for protein purity and integrity. Precision Plus Protein Standards Dual Color (BioRad), ranging from 10-250 kDa. was used as molecular weight markers. All samples (2.5 µg) were boiled for five minutes in Laemmli sample buffer before each run (79). Electrophoresis was performed for approximately 90 min in an ice bath at 25 mA constant current, and gels were stained using a Coomassie Blue Silver staining protocol (80).

### *In vitro* induction of Cnm amyloid formation

Six different protein variants were used in *in vitro* assays: Cnm (native, glycosylated), rCnm, rCBD, rAgA (amyloidogenic; positive control) and recombinant glucan binding protein C (rGpbC) (non-amyloidogenic; negative control). The purification of control proteins was described previously (15). Each protein was subjected to three different conditions that included stirring, stirring in the presence of the amyloid inhibitor Tannic Acid and unstirred (static). Mechanical agitation such as stirring promotes amyloid nucleation and is commonly used in studies investigating amyloidogenic properties of proteins (81). Two hundred microliters of each sample, which included PBS (1x, pH 7.4), Tannic Acid (100 µM, when present) and protein at 250 µg.mL^-1^, was placed in a 1.5 mL microcentrifuge tube with a 3×10 mm stir bar (except for the unstirred condition). All samples were incubated for five days at 4°C. Stirred samples, with or without Tannic Acid, were placed on magnetic stirrers at maximum intensity. Amyloid material obtained by this method was further used in the Thioflavin-T fluorescence assays, Transmission Electron Microscopy imaging and competitive binding assays.

### Thioflavin-T fluorescence assays

To test whether Cnm and its variant forms have amyloidogenic properties, we determined whether they uptake the amyloidophilic Thioflavin-T (ThT) as described elsewhere (15). Twenty-two microliters of a 20 µM ThT solution in PBS were added to stirred, stirred in the presence of Tannic Acid or unstirred samples prepared as described above, followed by incubation in the dark for 30 min at room temperature. Next, samples were transferred to fully dark 96 well plates and fluorescence was read in a Synergy H1 Hybrid Multi-Mode Reader (BioTek) with excitation at 440 nm and reading at 485 nm. A PBS + ThT blank was subtracted from each sample reading. To determine whether pH can influence amyloid formation, ThT assays were also performed as listed above but the pH was adjusted to pH 7.4 or 5.5. To account for potential pH interference of ThT incorporation, 20 µL of 20x PBS at pH 7.4 was added to the samples before the fluorescence readings. PBS + ThT blanks for each pH were subtracted from each sample reading. For all ThT Assays, results are displayed as relative fluorescence compared to stirred Cnm (at pH 7.4), in proportion to the molarity of each sample.

### Transmission Electron Microscopy (TEM) of amyloid material following treatment with proteinase K

Stirred samples of Cnm and its variants were treated with Proteinase K (ProK) at a 10:1 (amyloid:ProK) molar ratio to degrade residual protein monomers as previously described (48). Following incubation for 3 hours at 37°C, the reaction was stopped by addition of 2 mM phenylmethylsulfonyl fluoride (PMSF). After 5 min of incubation at room temperature, samples were centrifuged at 100,000 x g at 10°C for 30 min and the pellets were washed twice with distilled water and then resuspended in 100 µL of distilled water. Samples were visualized on SDS-PAGE to confirm absence of monomers (data not shown). Finally, samples were dried overnight, weighed on an analytical balance, and resuspended in distilled water to a concentration of 1 mg.mL^-1^. A 30 µL drop of each sample was placed under a 100-mesh formvar-carbon coated grid FCF100-Cu-UB (Electron Microscopy Sciences, Hatfield, PA) with 1% uranyl acetate. Imaging was done on a Hitachi H-7600 TEM (Hitachi High Technologies America, Schaumburg, IL) with digital images acquired at 80 keV using an AMT digital camera.

### Congo red birefringence assay of *S. mutans* biofilms

Evaluation of Congo red-induced birefringence of amyloid material within *S. mutans* biofilms was performed as described previously (15, 16). Briefly, overnight cultures of strains OMZ175, OMZ175*Δcnm* and OMZ175*ΔsrtA* grown in BHI broth were diluted 1:100 in Biofilm Media + 1% Glucose (BMG) (82). Two hundred microliters of the diluted cultures were added to 96 wells microtiter plates and incubated for 3 days at 37°C in a 5% CO_2_ atmosphere for biofilm formation. Spent media was then removed by aspiration and fresh BMG replaced daily until the biofilms were 7 days old. The wells were gently washed with 1x sterile PBS and biofilms were scraped off and centrifuged at 16,000 x g for 10 minutes. Pellets were resuspended in 10 μL of a Congo red solution, and incubated in microtubes for one hour at room temperature in the dark. CR solution was prepared by dissolving 2 g NaCl in 20 ml deionized H_2_O and by suspending 0.5 g CR in 80 ml 100% ethanol; these two solutions were combined, filtered and stored at room temperature as a stock solution. Samples were transferred to glass slides and visualized for birefringence as wet mounts using a Zeiss Scope A1 with a computer-controlled ProgRes C5 Jenoptik inverted camera equipped with cross-polarized filters. The number of birefringent foci were counted for each slide.

### *In vitro* and *ex vivo* collagen-binding competition assays

For *in vitro* competitive inhibition assays, wells of 96 wells plate were coated with 100 µL of 40 µg.mL^-1^ rat tail type I collagen in sterile PBS (Life Technologies, NY) at 4°C for 18 hours. Then, wells were gently washed three times with PBS and blocked with 200 µL of PBS containing 5% bovine serum albumin (m/v) for 1.5 hour at 37°C. The wells were washed again then incubated for 1 hour at 37°C with 50 µL of PBS (untreated), or PBS containing rCBD monomers or rCBD amyloid fibers at the indicated concentrations (0.2-10 µg). Following washing, 100 µL of bacterial suspensions of *S. mutans* OMZ175 grown in BHI to an optical density at 600 nm of ≈0.35 were added to quadruplicate wells of each experimental condition, and incubated for 3 hours at 37°C in 5% CO_2_. Following washing, wells were air dried and stained for 1 minute with 200 µL of 0.05% crystal violet (w/v), washed with PBS containing 0.01% Tween-20 (v/v) to remove excess unbound crystal violet, then treated with 200 µL of a 7% solution of acetic acid (v/v) to resuspend bound crystal violet. Absorbance was read at 575 nm on Synergy H1 Hybrid Multi-Mode Reader (BioTek). As a background control, absorbance was measured in wells containing collagen only (no bacteria) and subtracted from test values. Readings were normalized based on Colony Forming Unit (CFU) counts of the initial inocula plated on tryptic soy agar (TSA).

For *ex vivo* competitive inhibition assays we followed the protocol previously published by our lab (31) with some modifications. Twelve surgically extracted nonimpacted premolars that are typically discarded after extraction were obtained from the University of Florida’s dental clinic at the College of Dentistry and were cleaned of all soft tissue detritus and cut horizontally with a circular saw blade 17 mm from the crown, exposing a similar surface area of dentin on the root fragments. Prior to the assay, tooth fragments were rinsed clean by six rounds of 30 seconds of sonication in 4 mL of sterile water. Then, tooth fragments were disinfected for one hour in 70% ethanol and rehydrated overnight in 1x PBS. Surface disinfection was assessed by plating on TSA and incubation for 48 hours at 37°C and 5% CO_2_ (v/v). Each tooth fragment was then submersed in 1 mL of sterile clarified human pooled saliva for 30 minutes and incubated for 1 hour at 37°C in 500 µL of 1x PBS containing 20 µg of either rCBD monomer or rCBD purified amyloid fibers. Untreated tooth fragments served as a basis for comparison. Each sample group contained four replicates. Treated and untreated root sections were then submerged in 1 mL of a mid-log culture (OD_600_≈0.5) of OMZ175 in BHI broth for 5 minutes. Loosely bound cells were removed by three subsequent washes of each root fragment in 14 mL of sterile 1x PBS at 60 rpm. Strongly adherent cells were then removed by three rounds of sonication of 10 seconds each, in 1x PBS, with 30 seconds on ice between each round. Sonicants were plated on TSA and CFUs were enumerated after 48 hours of incubation at 37°C and 5% CO_2_ (v/v).

## Acknowledgements

The authors would like to thank Matthew Alzate, Ana Barran Berdon and Joshua Lovelace for technical assistance. This work was supported by NIH/NIDCR R01 DE021789 to L.J.B and R01 DE022559 to J.A. and J.A.L.

The authors declare no conflicts of interest

## Tables

**Table 1.**
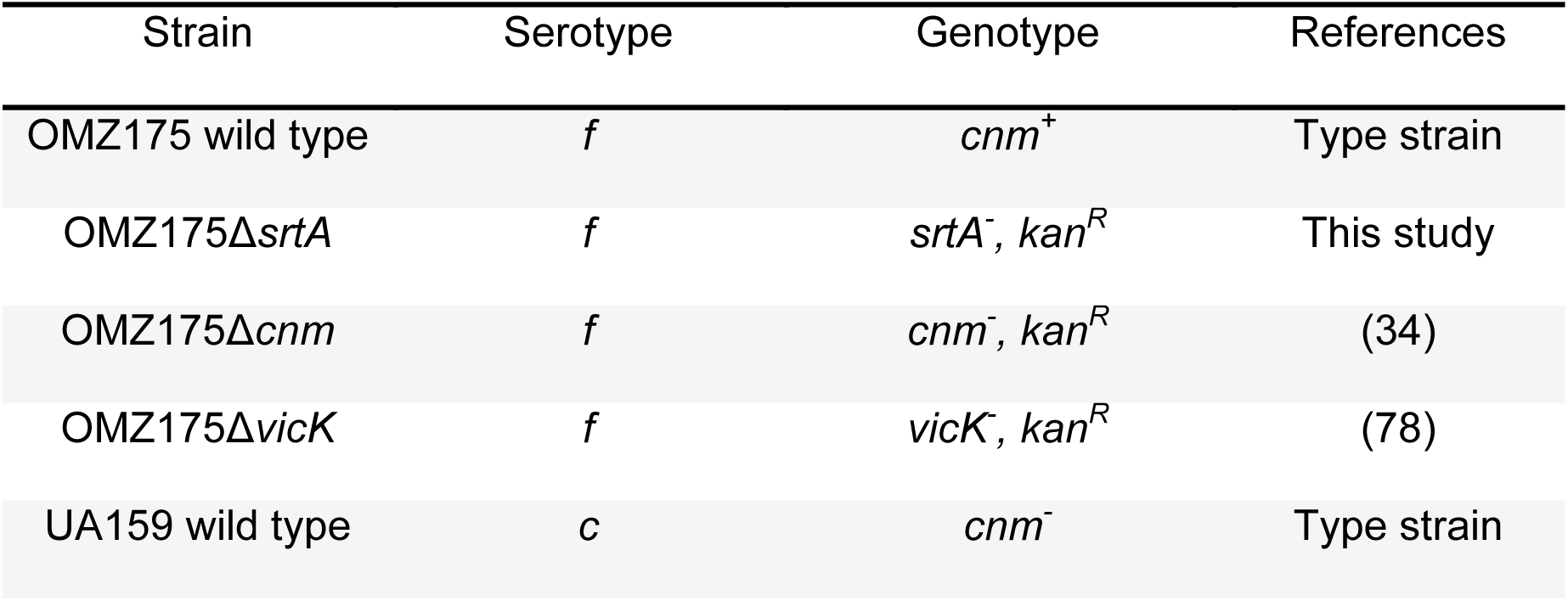
*Streptococcus mutans* strains used in this study.

**Table 2.** *Escherichia coli* strains used in this study.

| Strain | Genotype | References |
| --- | --- | --- |
| BL21 DE3 | F<sup>-</sup> *ompT hsdSB* (rB<sup>-</sup>mB<sup>-</sup>) *gal dcm* (λDE3) | Type strain |
| +pET-16b:rCBD | pET-16b carrying the CBD of Cnm (amino acids 32-319), *amp*<sup>R</sup> | (41) |
| +pET-16b:rCnm | pET-16b carrying full length Cnm (amino acids 32-545), *amp*<sup>R</sup> | (77) |

## References

1. Taglialegna A, Lasa I, Valle J. 2016. Amyloid Structures as Biofilm Matrix Scaffolds. J Bacteriol 198:2579–88.

2. Harrison RS, Sharpe PC, Singh Y, Fairlie DP. 2007. Amyloid peptides and proteins in review. Rev Physiol Biochem Pharmacol 159:1–77.

3. Shanmugam N, Baker MODG, Ball SR, Steain M, Pham CLL, Sunde M. 2019. Microbial functional amyloids serve diverse purposes for structure, adhesion and defence. Biophys Rev 11:287–302.

4. Greenwald J, Riek R. 2010. Biology of Amyloid: Structure, Function, and Regulation. Structure 18:1244–1260.

5. Howie AJ, Brewer DB. 2009. Optical properties of amyloid stained by Congo red: History and mechanisms. Micron 40:285–301.

6. Fowler DM, Koulov A V., Balch WE, Kelly JW. 2007. Functional amyloid - from bacteria to humans. Trends Biochem Sci 32:217–224.

7. Elliot MA, Karoonuthaisiri N, Huang J, Bibb MJ, Cohen SN, Kao CM, Buttner MJ. 2003. The chaplins: A family of hydrophobic cell-surface proteins involved in aerial mycelium formation in *Streptomyces coelicolor*. Genes Dev 17:1727–1740.

8. Seviour T, Hansen SH, Yang L, Yau YH, Wang VB, Stenvang MR, Christiansen G, Marsili E, Givskov M, Chen Y, Otzen DE, Nielsen PH, Geifman-Shochat S, Kjelleberg S, Dueholm MS. 2015. Functional amyloids keep quorum-sensing molecules in check. J Biol Chem 290:6457–6469.

9. DePas WH, Chapman MR. 2012. Microbial manipulation of the amyloid fold. Res Microbiol 163:592–606.

10. Erskine E, MacPhee CE, Stanley-Wall NR. 2018. Functional Amyloid and Other Protein Fibers in the Biofilm Matrix. J Mol Biol 430:3642–3656.

11. Jain N, Chapman MR. 2019. Bacterial functional amyloids: Order from disorder. Biochim Biophys Acta - Proteins Proteomics 1867:954–960.

12. Salinas N, Povolotsky TL, Landau M, Kolodkin-Gal I. 2020. Emerging Roles of Functional Bacterial Amyloids in Gene Regulation, Toxicity, and Immunomodulation. Microbiol Mol Biol Rev 85:1–17.

13. Andreasen M, Meisl G, Taylor JD, Michaels TCT, Levin A, Otzen DE, Chapman MR, Dobson CM, Matthews SJ, Knowles TPJ. 2019. Physical determinants of amyloid assembly in biofilm formation. MBio 10:1–12.

14. Cámara-Almirón J, Caro-Astorga J, de Vicente A, Romero D. 2018. Beyond the expected: the structural and functional diversity of bacterial amyloids. Crit Rev Microbiol 44:653–666.

15. Besingi RN, Wenderska IB, Senadheera DB, Cvitkovitch DG, Long JR, Wen ZT, Brady LJ. 2017. Functional amyloids in *Streptococcus mutans*, their use as targets of biofilm inhibition and initial characterization of SMU_63c. Microbiol (United Kingdom) 163:488–501.

16. Oli MW, Otoo HN, Crowley PJ, Heim KP, Nascimento MM, Ramsook CB, Lipke PN, Brady LJ. 2012. Functional amyloid formation by *Streptococcus mutans*. Microbiology 158:2903–2916.

17. Taglialegna A, Matilla-Cuenca L, Dorado-Morales P, Navarro S, Ventura S, Garnett JA, Lasa I, Valle J. 2020. The biofilm-associated surface protein Esp of *Enterococcus faecalis* forms amyloid-like fibers. Biofilms and Microbiomes 6:1–12.

18. Schwartz K, Syed AK, Stephenson RE, Rickard AH, Boles BR. 2012. Functional amyloids composed of phenol soluble modulins stabilize Staphylococcus aureus biofilms. PLoS Pathog 8:1–11.

19. Baldry M, Bojer MS, Najarzadeh Z, Vestergaard M, Meyer RL, Otzen DE, Ingmer H. 2020. Phenol-Soluble Modulins Modulate Persister Cell Formation in *Staphylococcus aureus*. Front Microbiol 11:1–7.

20. Chai L, Romero D, Kayatekin C, Akabayov B, Vlamakis H, Losick R, Kolter R. 2013. Isolation, characterization, and aggregation of a structured bacterial matrix precursor. J Biol Chem 288:17559–17568.

21. Chen D, Cao Y, Yu L, Tao Y, Zhou Y, Zhi Q, Lin H. 2019. Characteristics and influencing factors of amyloid fibers in *S. mutan*s biofilm. AMB Express 9:1–9.

22. Bowen WH, Burne RA, Wu H, Koo H. 2018. Oral Biofilms: Pathogens, Matrix, and Polymicrobial Interactions in Microenvironments. Trends Microbiol 26:229–242.

23. Lemos JA, Palmer SR, Zeng L, Wen ZT, Kajfasz JK, Freires IA, Abranches J, Brady LJ. 2019. The Biology of *Streptococcus mutans*. Microbiol Spectr 7:1–18.

24. Lemos JA, Quivey RG, Koo H, Abranches J. 2013. *Streptococcus mutans*: a new Gram-positive paradigm? Microbiology 159:436–445.

25. Zhu L, Kreth J, Cross SE, Gimzewski JK, Shi W, Qi F. 2006. Functional characterization of cell-wall-associated protein WapA in *Streptococcus mutans*. Microbiology 152:2395–2404.

26. Sullan RMA, Li JK, Crowley PJ, Brady LJ, Dufrêne YF. 2015. Binding forces of *Streptococcus mutans* P1 adhesin. ACS Nano 9:1448–1460.

27. Avilés-Reyes A, Miller JHH, Lemos JAA, Abranches J. 2017. The collagen-binding proteins of *Streptococcus mutans* and related streptococci. Mol Oral Microbiol 32:89–106.

28. Nakano K, Nomura R, Taniguchi N, Lapirattanakul J, Kojima A, Naka S, Senawongse P, Srisatjaluk R, Grönroos L, Alaluusua S, Matsumoto M, Ooshima T. 2010. Molecular characterization of *Streptococcus mutans* strains containing the cnm gene encoding a collagen-binding adhesin. Arch Oral Biol 55:34–39.

29. Nomura R, Nakano K, Taniguchi N, Lapirattanakul J, Nemoto H, Grönroos L, Alaluusua S, Ooshima T. 2009. Molecular and clinical analyses of the gene encoding the collagen-binding adhesin of *Streptococcus mutans*. J Med Microbiol 58:469–475.

30. Nakano K, Lapirattanakul J, Nomura R, Nemoto H, Alaluusua S, Grönroos L, Vaara M, Hamada S, Ooshima T, Nakagawa I. 2007. *Streptococcus mutans* clonal variation revealed by multilocus sequence typing. J Clin Microbiol 45:2616–2625.

31. Miller JH, Avilés-Reyes A, Scott-Anne K, Gregoire S, Watson GE, Sampson E, Progulske-Fox A, Koo H, Bowen WH, Lemos JA, Abranches J. 2015. The collagen binding protein Cnm contributes to oral colonization and cariogenicity of *Streptococcus mutans* OMZ175. Infect Immun 83:2001–2010.

32. Nomura R, Naka S, Nemoto H, Otsugu M, Nakamura S, Ooshima T, Nakano K. 2013. Potential high virulence for infective endocarditis in *Streptococcus mutans* strains with collagen-binding proteins but lacking PA expression. Arch Oral Biol 58:1627–1634.

33. Arora S, Gordon J, Hook M. 2021. Collagen Binding Proteins of Gram-Positive Pathogens. Front Microbiol 12:1–16.

34. Abranches J, Miller JH, Martinez AR, Simpson-Haidaris PJ, Burne RA, Lemos JA. 2011. The collagen-binding protein Cnm is required for *Streptococcus mutans* adherence to and intracellular invasion of human coronary artery endothelial cells. Infect Immun 79:2277–2284.

35. Inenaga C, Hokamura K, Nakano K, Nomura R, Naka S, Ohashi T, Ooshima T, Kuriyama N, Hamasaki T, Wada K, Umemura K, Tanaka T. 2018. A Potential New Risk Factor for Stroke: *Streptococcus mutans* With Collagen-Binding Protein. World Neurosurg 113:e77–e81.

36. Nomura R, Otsugu M, Hamada M, Matayoshi S, Teramoto N, Iwashita N, Naka S, Matsumoto-Nakano M, Nakano K. 2020. Potential involvement of *Streptococcus mutans* possessing collagen binding protein Cnm in infective endocarditis. Sci Rep 10:1–14.

37. Miyatani F, Kuriyama N, Watanabe I, Nomura R, Nakano K, Matsui D, Ozaki E, Koyama T, Nishigaki M, Yamamoto T, Mizuno T, Tamura A, Akazawa K, Takada A, Takeda K, Yamada K, Nakagawa M, Ihara M, Kanamura N, Friedland RP, Watanabe Y. 2015. Relationship between Cnm-positive *Streptococcus mutans* and cerebral microbleeds in humans. Oral Dis 21:886–893.

38. Hosoki S, Saito S, Tonomura S, Ishiyama H, Yoshimoto T, Ikeda S, Ikenouchi H, Yamamoto Y, Hattori Y, Miwa K, Friedland RP, Carare RO, Nakahara J, Suzuki N, Koga M, Toyoda K, Nomura R, Nakano K, Takegami M, Ihara M. 2020. Oral Carriage of *Streptococcus mutans* Harboring the *cnm* Gene Relates to an Increased Incidence of Cerebral Microbleeds. Stroke 3632–3639.

39. Freires IA, Avilés-Reyes A, Kitten T, Simpson-Haidaris PJ, Swartz M, Knight PA, Rosalen PL, Lemos JA, Abranches J. 2017. Heterologous expression of *Streptococcus mutans* Cnm in *Lactococcus lactis* promotes intracellular invasion, adhesion to human cardiac tissues and virulence. Virulence 8:18–29.

40. Garcia BA, Acosta NC, Tomar SL, Roesch LFW, Lemos JA, Mugayar LRF, Abranches J. 2021. Association of *Candida albicans* and Cbp+ *Streptococcus mutans* with early childhood caries recurrence. Sci Rep 11:1–11.

41. Avilés-Reyes A, Freires IA, Besingi R, Purushotham S, Deivanayagam C, Brady LJ, Abranches J, Lemos JA. 2018. Characterization of the pgf operon involved in the posttranslational modification of *Streptococcus mutans* surface proteins. Sci Rep 8:1–12.

42. Avilés-Reyes A, Miller JH, Simpson-Haidaris PJ, Hagen FK, Abranches J, Lemos JA. 2014. Modification of S*treptococcus mutans* Cnm by PgfS contributes to adhesion, endothelial cell invasion, and virulence. J Bacteriol 196:2789–2797.

43. Ono K, Hasegawa K, Naiki H, Yamada M. 2004. Anti-amyloidogenic activity of tannic acid and its activity to destabilize Alzheimer’s β-amyloid fibrils *in vitro*. Biochim Biophys Acta - Mol Basis Dis 1690:193–202.

44. Xue C, Lin TY, Chang D, Guo Z. 2017. Thioflavin T as an amyloid dye: fibril quantification, optimal concentration and effect on aggregation. R Soc Open Sci 4:1–12.

45. Van Gerven N, Van der Verren SE, Reiter DM, Remaut H. 2018. The Role of Functional Amyloids in Bacterial Virulence. J Mol Biol 430:3657–3684.

46. Lassé M, Ulluwishewa D, Healy J, Thompson D, Miller A, Roy N, Chitcholtan K, Gerrard JA. 2016. Evaluation of protease resistance and toxicity of amyloid-like food fibrils from whey, soy, kidney bean, and egg white. Food Chem 192:491–498.

47. Kushnirov V V., Dergalev AA, Alexandrov AI. 2020. Proteinase K resistant cores of prions and amyloids. Prion 14:11–19.

48. Barran-Berdon AL, Ocampo S, Haider M, Morales-Aparicio J, Ottenberg G, Kendall A, Yarmola E, Mishra S, Long JR, Hagen SJ, Stubbs G, Brady LJ. 2020. Enhanced purification coupled with biophysical analyses shows cross-β structure as a core building block for *Streptococcus mutans* functional amyloids. Sci Rep 10:1–11.

49. Howie AJ, Brewer DB, Howell D, Jones AP. 2008. Physical basis of colors seen in Congo red-stained amyloid in polarized light. Lab Investig 88:232–242.

50. Goldberg M. 2011. Dentin structure composition and mineralization. Front Biosci 3:711–735.

51. Lo Giudice G, Cutroneo G, Centofanti A, Artemisia A, Bramanti E, Militi A, Rizzo G, Favaloro A, Irrera A, Lo Giudice R, Cicciù M. 2015. Dentin Morphology of Root Canal Surface: A Quantitative Evaluation Based on a Scanning Electronic Microscopy Study. Biomed Res Int 2015:1–7.

52. Wösten HAB, de Vocht ML. 2000. Hydrophobins, the fungal coat unravelled. Biochim Biophys Acta - Rev Biomembr 1469:79–86.

53. Nelson R, Sawaya MR, Balbirnie M, Madsen A, Riekel C, Grothe R, Eisenberg D. 2005. Structure of the cross-β spine of amyloid-like fibrils. Nature 435:773–778.

54. Fowler DM, Koulov A V., Alory-Jost C, Marks MS, Balch WE, Kelly JW. 2005. Functional Amyloid Formation within Mammalian Tissue. PLoS Biol 4:100–107.

55. Maji SK, Perrin MH, Sawaya MR, Jessberger S, Vadodaria K, Rissman RA, Singru PS, Nilsson KPR, Simon R, Schubert D, Eisenberg D, Rivier J, Sawchenko P, Vale W, Riek R. 2009. Functional Amyloids As Natural Storage of Peptide Hormones in Pituitary Secretory Granules. Science (80-) 325:328–332.

56. Boix CP, Lopez-Font I, Cuchillo-Ibañez I, Sáez-Valero J. 2020. Amyloid precursor protein glycosylation is altered in the brain of patients with Alzheimer’s disease. Alzheimers Res Ther 12:96.

57. Schedin-Weiss S, Winblad B, Tjernberg LO. 2014. The role of protein glycosylation in Alzheimer disease. FEBS J 281:46–62.

58. Lin Y, Reino C, Carrera J, Pérez J, van Loosdrecht MCM. 2018. Glycosylated amyloid- like proteins in the structural extracellular polymers of aerobic granular sludge enriched with ammonium- oxidizing bacteria. Microbiologyopen 7:1–13.

59. Esberg A, Sheng N, Mårell L, Claesson R, Persson K, Borén T, Strömberg N. 2017. Streptococcus mutans adhesin biotypes that match and predict individual caries development. EBioMedicine 24:205–215.

60. Sharma A, Parashar D, Satyanarayana T. 2016. Acidophilic Microbes: Biology and Applications, p. 215–241. In Rampelotto, PH (ed.), Biotechnology of Extremophiles, 1st ed. Springer International Publishing.

61. Paredes-Rosan CA, Valencia DE, Barazorda-Ccahuana HL, Aguilar-Pineda JA, Gómez B. 2020. Amyloid beta oligomers: how pH influences over trimer and pentamer structures? J Mol Model 26:1–8.

62. Su Y, Chang P-T. 2001. Acidic pH promotes the formation of toxic fibrils from β-amyloid peptide. Brain Res 893:287–291.

63. Taglialegna A, Navarro S, Ventura S, Garnett JA, Matthews S, Penades JR, Lasa I, Valle J. 2016. Staphylococcal Bap Proteins Build Amyloid Scaffold Biofilm Matrices in Response to Environmental Signals. PLoS Pathog 12:1–34.

64. Lévesque CM, Voronejskaia E, Huang YCC, Mair RW, Ellen RP, Cvitkovitch DG. 2005. Involvement of sortase anchoring of cell wall proteins in biofilm formation by *Streptococcus mutans*. Infect Immun 73:3773–3777.

65. Sawyer EB, Claessen D, Haas M, Hurgobin B, Gras SL. 2011. The Assembly of Individual Chaplin Peptides from *Streptomyces coelicolor* into Functional Amyloid Fibrils. PLoS One 6:1–12.

66. Palmer SR, Miller JH, Abranches J, Zeng L, Lefebure T, Richards VP, Lemos JA, Stanhope MJ, Burne RA. 2013. Phenotypic Heterogeneity of Genomically-Diverse Isolates of *Streptococcus mutans*. PLoS One 8:1–17.

67. Shammas SL, Knowles TPJ, Baldwin AJ, MacPhee CE, Welland ME, Dobson CM, Devlin GL. 2011. Perturbation of the stability of amyloid fibrils through alteration of electrostatic interactions. Biophys J 100:2783–2791.

68. Otto M. 2013. Staphylococcal infections: Mechanisms of biofilm maturation and detachment as critical determinants of pathogenicity. Annu Rev Med 64:175–188.

69. Harper JD, Lansbury PT. 1997. Models of amyloid seeding in Alzheimer’s disease and scrapie: Mechanistic truths and physiological consequences of the time-dependent solubility of amyloid proteins. Annu Rev Biochem 66:385–407.

70. Álvarez-Mena A, Cámara-Almirón J, de Vicente A, Romero D. 2020. Multifunctional amyloids in the biology of Gram-positive bacteria. Microorganisms 8:1–20.

71. Lau PCY, Sung CK, Lee JH, Morrison DA, Cvitkovitch DG. 2002. PCR ligation mutagenesis in transformable streptococci: Application and efficiency. J Microbiol Methods 49:193–205.

72. Kremer BHA, van der Kraan M, Crowley PJ, Hamilton IR, Brady LJ, Bleiweis AS. 2001. Characterization of the sat Operon in *Streptococcus mutans*: Evidence for a Role of Ffh in Acid Tolerance. J Bacteriol 183:2543–2552.

73. Tsolis AC, Papandreou NC, Iconomidou VA, Hamodrakas SJ. 2013. A Consensus Method for the Prediction of “Aggregation-Prone” Peptides in Globular Proteins. PLoS One 8:1–6.

74. Waterhouse A, Bertoni M, Bienert S, Studer G, Tauriello G, Gumienny R, Heer FT, de Beer TAP, Rempfer C, Bordoli L, Lepore R, Schwede T. 2018. SWISS-MODEL: homology modelling of protein structures and complexes. Nucleic Acids Res 46:W296–W303.

75. Baek M, Park T, Heo L, Park C, Seok C. 2017. GalaxyHomomer: a web server for protein homo-oligomer structure prediction from a monomer sequence or structure. Nucleic Acids Res 45:W320–W324.

76. Piovesan D, Necci M, Escobedo N, Monzon AM, Hatos A, Mičetić I, Quaglia F, Paladin L, Ramasamy P, Dosztányi Z, Vranken WF, Davey NE, Parisi G, Fuxreiter M, Tosatto SCE. 2021. MobiDB: intrinsically disordered proteins in 2021. Nucleic Acids Res 49:D361–D367.

77. Avilés-Reyes A, Miller JH, Simpson-Haidaris PJ, Lemos JA, Abranches J. 2014. Cnm is a major virulence factor of invasive *Streptococcus mutans* and part of a conserved three-gene locus. Mol Oral Microbiol 29:11–23.

78. Alves LA, Ganguly T, Mattos-Graner RO, Kajfasz J, Harth-Chu EN, Lemos JA, Abranches J, Araújo Alves L, Ganguly T, Mattos-Graner RO, Kajfasz J, Harth-Chu EN, Lemos JA, Abranches J. 2018. CovR and VicRKX regulate transcription of the collagen binding protein Cnm of *Streptococcus mutans*. J Bacteriol 200:1–14.

79. Laemmli UK. 1970. Cleavage of structural proteins during the assembly of the head of bacteriophage T4. Nature 227:680–685.

80. Candiano G, Bruschi M, Musante L, Santucci L, Ghiggeri GM, Carnemolla B, Orecchia P, Zardi L, Righetti PG. 2004. Blue silver: A very sensitive colloidal Coomassie G-250 staining for proteome analysis. Electrophoresis 25:1327–1333.

81. Bäcklund FG, Pallbo J, Solin N, Blacklow S. 2016. Controlling amyloid fibril formation by partial stirring. Biopolymers 105:249–259.

82. Lemos JA, Abranches J, Koo H, Marquis RE, Burne RA. 2010. Protocols to study the physiology of oral biofilms. Methods Mol Biol 666:87–102.

